# Brain network motifs are markers of loss and recovery of consciousness

**DOI:** 10.1101/2020.03.16.993659

**Authors:** Catherine Duclos, Danielle Nadin, Yacine Mahdid, Vijay Tarnal, Paul Picton, Giancarlo Vanini, Goodarz Golmirzaie, Ellen Janke, Michael S. Avidan, Max B. Kelz, George A. Mashour, Stefanie Blain-Moraes

**Author notes:** **Corresponding Author:** Stefanie Blain-Moraes, Ph.D. Montreal General Hospital, Room L11-132, 1650 Cedar Ave, L11-132 Montréal, Québec, H3G 1A4 Canada. These authors contributed equally to the work (Catherine Duclos, Danielle Nadin).

## Abstract

Motifs are patterns of inter-connections between nodes of a network, and have been investigated as building blocks of directed networks. This study explored the re-organization of 3-node motifs during loss and recovery of consciousness. Nine healthy subjects underwent a 3-hour anesthetic protocol while 128-channel electroencephalography (EEG) was recorded. In the alpha (8–13 Hz) band, five-minute epochs of EEG were extracted for: baseline; induction; unconscious; 30-, 10- and 5-minutes pre-recovery of responsiveness; 30- and 180-minutes post-recovery of responsiveness. We constructed a functional brain network using the weighted and directed phase lag index, on which we calculated the frequency and topology of 3-node motifs. Three motifs (motifs 1, 2 and 5) were significantly present across participants and epochs, when compared to random networks (p<0.05). The topology of motifs 1 and 5 changed significantly between responsive and unresponsive epochs (p<0.01). Motif 1 was constituted by long-range chain-like connections, while motif 5 was constituted by short-range, loop-like connections. Our results suggest that anesthetic-induced unconsciousness is associated with a topological re-organization of network motifs. As motif topological re-organization may precede (motif 5) or accompany (motif 1) the return of responsiveness, motifs could contribute to the understanding of the neural correlates of consciousness.

## Introduction

The field of network neuroscience has yielded powerful insight into the way the brain is structurally and functionally connected. The application of graph theoretical analysis to neuroimaging techniques has provided evidence that the brain has features of a complex small-world network, with functional modules and densely connected hubs [1,2]. The development of measures quantifying properties of the brain network has led to a better understanding of the anatomical and functional architecture that drives various brain states, including anesthetic-induced unconsciousness. Both electroencephalography (EEG) and functional magnetic resonance imaging (fMRI) studies have shown that propofol-, sevoflurane-, and ketamine- induced unconsciousness induce a functional disconnection of anterior and posterior regions of the cortex [3–8], while propofol and sevoflurane also induce an anteriorization of alpha power from the occipital to the frontal cortex [9–11]. Graph theoretical analysis has demonstrated that global and topological properties of brain networks are altered during anesthetic-induced unconsciousness [12–16]. While these advances have led to a clearer understanding of how anesthetic-induced unconsciousness creates conditions incompatible with information processing and transfer [17], the majority of these measures describe macro-scale global network properties, using a single-number value, rather than meso- or micro-scale node-based changes in brain functioning.

Motifs are patterns of inter-connections between the nodes of complex networks, with a probability of occurrence that is significantly higher than those in randomized networks [18]. As such, functional motifs have been investigated as the basic building blocks of directed networks [18–19]. The distribution of motifs across a brain network is organized to support information integration and segregation – key properties associated with consciousness [20–22]. Shin *et al.* showed that motif frequency changes across anesthetic induction, maintenance and recovery, and that some motifs are state-specific [23]. Conversely, Kafashan *et al.* showed that sevoflurane-induced unconsciousness disrupts motifs of weaker correlation strength within and between resting-state networks, but preserves motifs with higher correlation strength [24]. These findings reveal changes in the building blocks of a network, on a nodal level, across states of consciousness, and suggest that motifs may reflect the granular network alterations of anesthesia-induced unconsciousness. However, the temporal and topological relationship between motifs and emergence from anesthetic-induced unconsciousness has yet to be established.

This exploratory study tested the hypothesis that loss and recovery of anesthetic-induced unconsciousness causes a re-organization of network motifs. More specifically, we aimed to determine the temporal association between network motifs and the behavioral return of responsiveness. Nine healthy adults underwent a controlled, 3-hour anesthetic protocol, during which high-density EEG was acquired. In the alpha frequency band (8–13 Hz), five-minute epochs of EEG were analyzed across the anesthetic protocol, for: baseline; induction; unconsciousness; 30-, 10, and 5-minutes prior to recovery of responsiveness; as well as 30- and 180-minutes post-recovery of responsiveness (Fig. 1). We hypothesized that different global states of consciousness [25] have a characteristic motif frequency and topology, and that motifs can effectively distinguish consciousness from anesthetic-induced unconsciousness. Given that motifs may constitute the building blocks of functional networks, we also hypothesized that the motifs associated with consciousness would be disrupted upon anesthetic-induced unconsciousness and would recover prior to the return of behavioral responsiveness. As nodal measures of functional brain networks, we hypothesized that motif frequency and topology would be useful measures to complement global network properties in the study of the neural correlates of consciousness.

**Figure 1.**
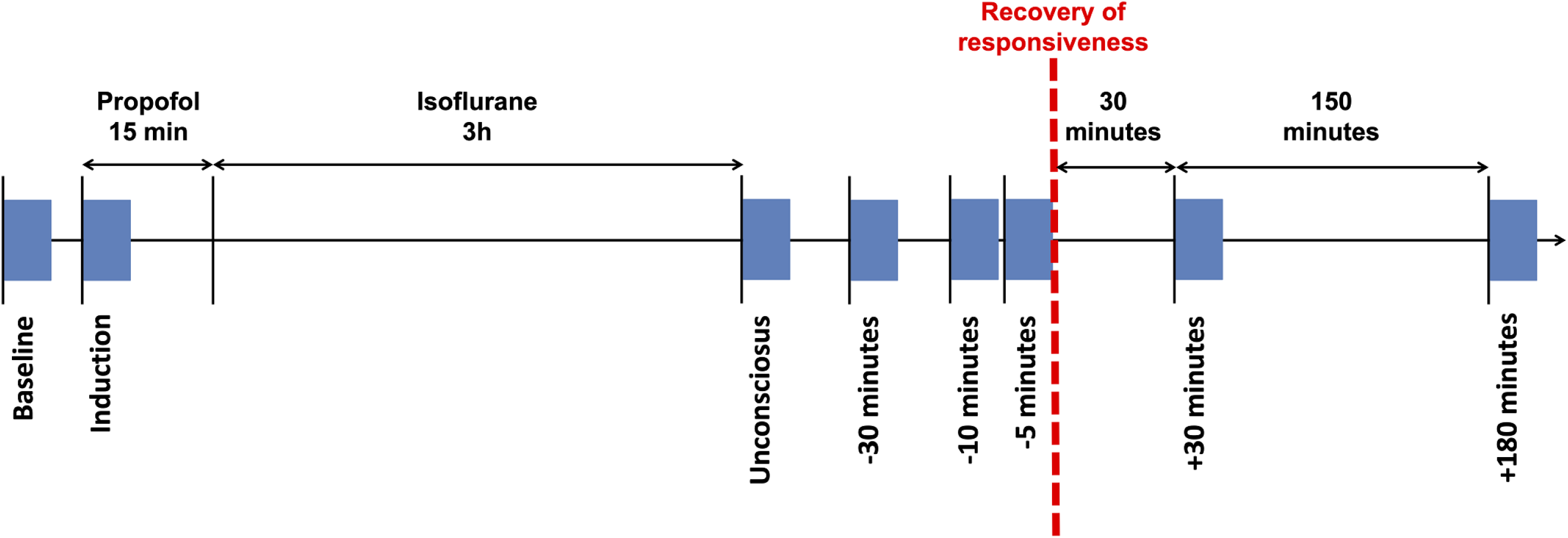
Experimental design and timeline. Timeline of anesthetic protocol and EEG data epochs. Participants received a stepwise increasing infusion rate of propofol for 15 minutes: 100 mcg/kg/min × 5 minutes increasing to 200 mcg/kg/min × 5 minutes, and then to 300 mcg/kg/min × 5 minutes. Participant then received 1.3 age-adjusted minimum alveolar concentration inhaled isoflurane anesthesia for 3 hours. Blue rectangles represent the eight 5-min EEG epochs during which network properties and motifs were calculated: 1) baseline; 2) induction; 3) unconscious; 4) 30 min prior to recovery of responsiveness; 5) 10 min prior to recovery of responsiveness; 6) 5 min prior to recovery of responsiveness; 7) 30 min post-recovery of responsiveness; and 8) 180 min post-recovery of responsiveness.

## Results

Nine healthy participants underwent a 3-hour anesthetic protocol at surgical levels while 128-channel EEG was recorded. The anesthetic protocol was comprised of a 15-minute propofol induction followed by a 3-hour period of isoflurane inhalation at 1.3 minimum alveolar concentration (MAC). Five-minute epochs of EEG were extracted for the following time points: 1) baseline; 2) induction; 3) unconscious; 4) 30 minutes prior to recovery of responsiveness (ROR); 5) 10 minutes prior to ROR; 6) 5 minutes prior to ROR; 7) 30 minutes post-ROR; and 8) 180 minutes post-ROR (Fig. 1). Half of these epochs (1,2,7,8) therefore represented periods of behavioral responsiveness and half (3-6) represented periods of behavioral unresponsiveness.

### Motifs are present across all states of consciousness, but their frequency cannot distinguish between responsive and unresponsive states

The total frequency of occurrence of the five unidirectional 3-node motifs (Fig. 2) was compared against 100 null networks to assess motif significance. Across all eight epochs, the frequency of motifs 1, 2 and 5 was significantly higher than in null networks, though this significance varied across epochs and participants (Fig. 3). Motif 1 was significantly present across epochs and participants (alpha: 98.6%), followed by motif 2 (95.8%) and motif 5 (91.7%). As motifs 3 and 4 did not appear significantly more frequently than in null networks, they were removed from subsequent analyses. The frequency of significant motifs (i.e. 1, 2 and 5) was compared across all epochs. The null hypothesis (H_0_) that there was no significant difference in motif frequency across responsive and unresponsive epochs could not be rejected with the Friedman test (*p* > 0.05). A Bayesian repeated-measures ANOVA yielded anecdotal evidence for H_0_ for motif 1 (*BF*_10_ = 0.879); moderate evidence for motif 2 (*BF*_10_ = 0.212); and anecdotal evidence for H_0_ for Motif 5 (*BF*_10_ = 0.908), suggesting that there was insufficient evidence to conclude that the total number of motifs changed across the experiment.

**Figure 2.**
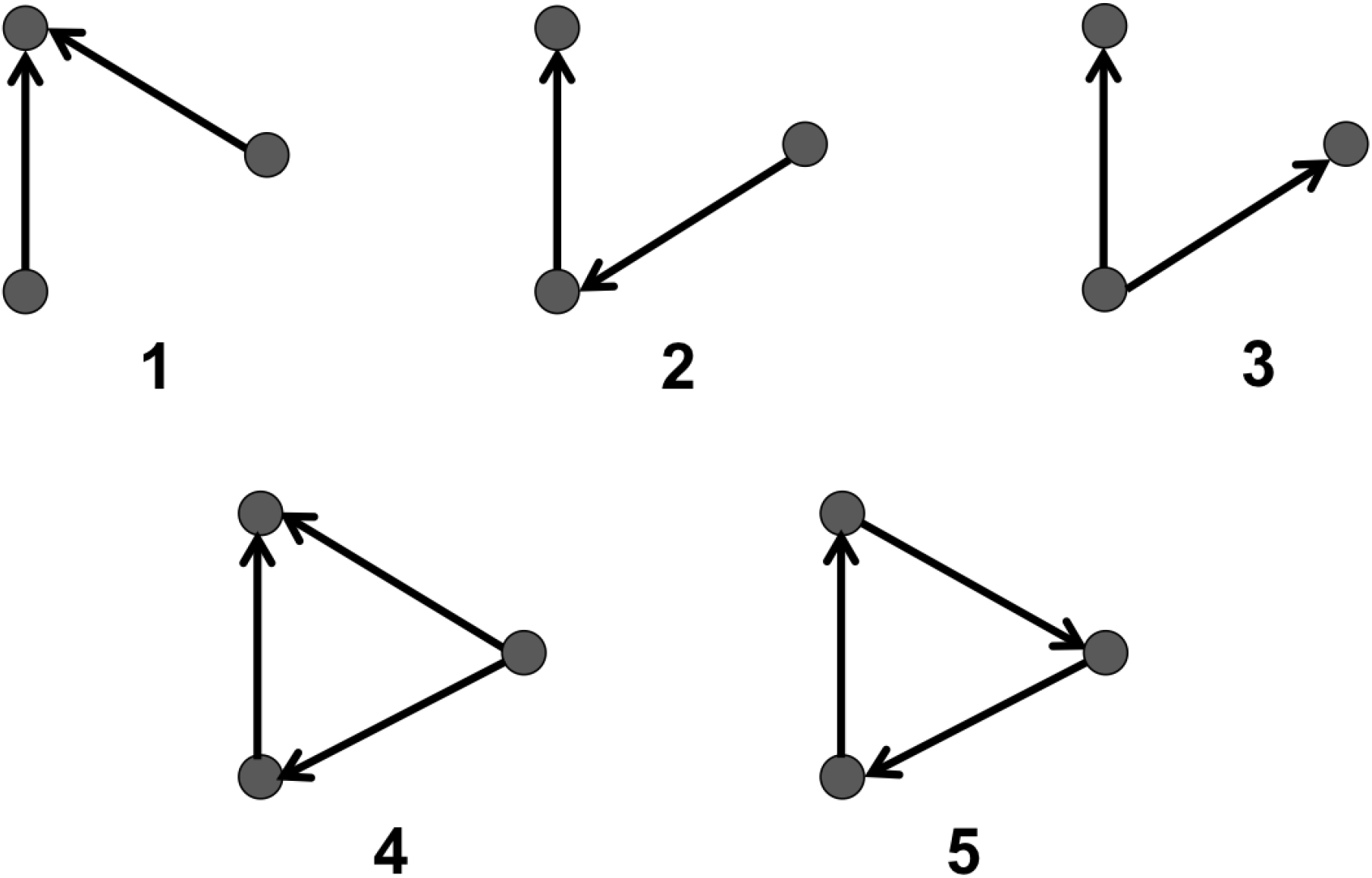
Possible 3-node network motifs. A 3-node motif represents a specific pattern of interconnection between three electrodes Five 3-node motifs can exist when unidirectional connections are assessed. These five 3-node motifs are assessed in this study. Dark circles represent nodes (i.e. individual electrodes), while black lines represent the edges linking the nodes. Arrows indicate the direction of phase-lead relationship.

**Figure 3.**
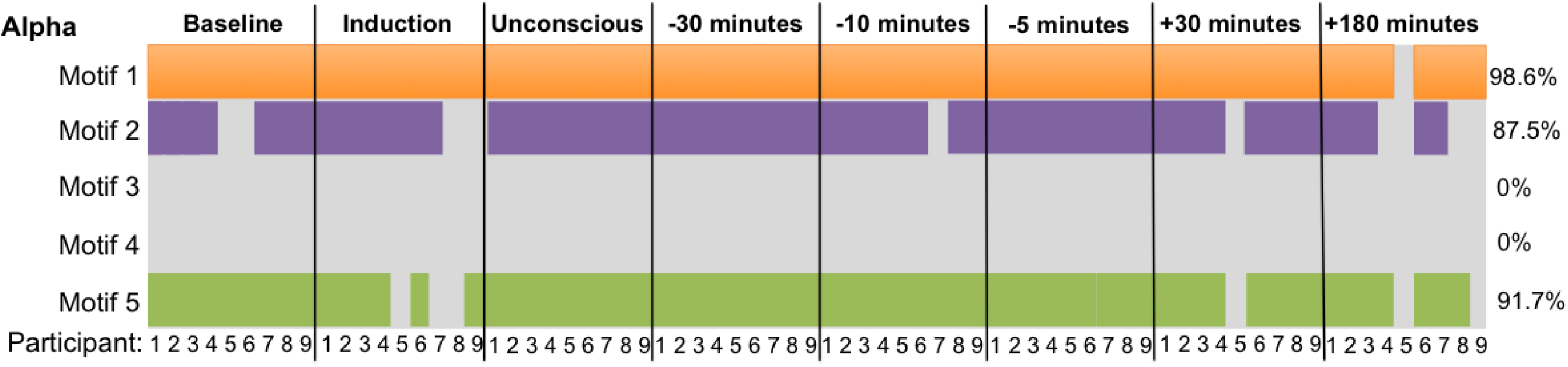
Motif significance. For each motif, the total aggregate frequency of motifs in the network was calculated by summing across all channels. A network Z-score was calculated per motif against the distribution of the total motifs in the 100 null networks. Motifs that were statistically significance per participant and time point are represented by orange (motif 1), purple (motif 2) and green (motif 5) rectangles. Motifs not statistically significant are left grey. Each time point has the potential to be significant in all nine participants, but motif significance varied across time points and participants. Percentages in the right column represent the frequency of motif significance. A frequency of 98.6% in motif significance indicates that this motif was significant throughout all participants and time points, except one participant, at one time point.

### Reconfiguration of motif topology marks changes in states of consciousness

On average across all participants, in the alpha band, nodes that participated in motifs 1 and 5 during conscious wakefulness were consolidated into circumscribed brain areas, with motif 1 dominant in central regions (Fig. 4A) and motif 5 dominant in posterior and peripheral regions (Fig. 4C). Nodes that participated in motif 2 were scattered across the brain and did not cluster into a single spatial pattern. The topologic distribution of motif 1 changed significantly from baseline (*χ*^2^(7) = 27.887, p < 0.001, W = 0.664) across all four unresponsive epochs: unconscious (p = 0.002); 30-minutes pre-ROR (p = 0.0065);10-minutes pre-ROR (p = 0.001); and 5-minutes pre-ROR (p = 0.00575) (Fig. 4A). The topologic distribution of motif 5 changed significantly from baseline (*χ*^2^(7) = 20.400, p = 0.005, W = 0.529) during the unconscious epoch (p = 0.0014) and 30-min pre-ROR (p = 0.0021) (Fig. 4C). The topologic distribution of motif 2 did not differ significantly from baseline at any epoch. While individual patterns of motif topology and re-organization varied slightly from this average pattern (Fig. 4E), topological reorganization was more visually apparent in individual subjects than on average for 7 of the 9 participants (see Supplementary Material Figure 1 for examples of individual figures). Topological re-organization in the theta band was similar to that observed in the alpha band, while there was no clear re-organization in the delta or beta bands (see Supplementary Material Figures 2, 3 and 4 for delta, theta, and beta results, respectively).

**Figure 4.**
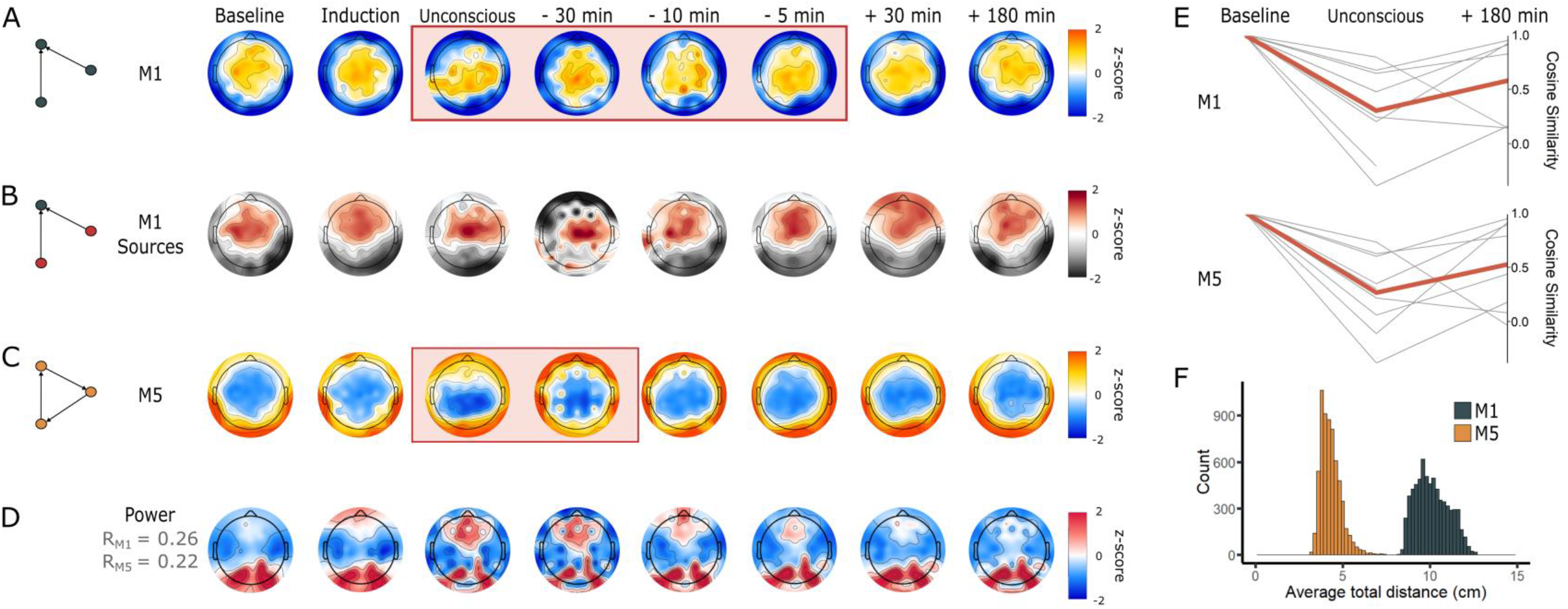
Motif topology across states of consciousness. A and C) Topology of the frequency of participation of nodes in alpha motif 1 (M1) and motif 5 (M5) across the eight analysis epochs. Orange rectangles highlight the epochs that are significantly distinct from Baseline (*p* < 0.05). The colormap represents the Z-score of the normalized frequency of motif participation for each electrode. B) Topographic maps depicting the distribution of nodes acting as sources of information flow (i.e. nodes that have a phase-lead relationship with regards to the other nodes within the motif), in every instance of motif 1 within the EEG network, across the eight time points of the anesthetic protocol. The colormap represents the Z-score of the total source score of each node. D) Alpha power topographic maps across the eight analysis epochs. There is a significant, small-to-medium positive correlation between alpha power the frequency of motif 1 (R = 0.26, p < 0.001) and motif 5 (R = 0.22, p < 0.001). E) Differences in cosine similarity values between Baseline, Unconsciousness, and 180 minutes post-recovery of consciousness. Grey lines depict single participants, while the red line depicts average cosine similarity value across all participants. F) Across participants and epochs, nodes participating in motif 1 form longer-range connections than those participating in motif 5.

### Topologic distribution of source nodes and alpha power

Using the dPLI, we assessed the lead-lag relationships in every motif to identify which nodes were origins of information flow (i.e. “sources”), and which were destinations of information (i.e. “sinks”). This source node analysis was only conducted for motif 1, given that all nodes in motif 5 are both the source and destination of information flow. Source nodes in motif 1 were located predominantly in anterior regions during baseline consciousness, and shifted towards dominance in central regions during the unconscious, 30- and 10-minutes pre-ROR epochs (Fig. 4B). Sources returned to anterior dominance upon recovery of responsiveness.

We investigated the association between the topologic distribution of alpha power and the changes in anterior-posterior dominance observed in motifs across states of consciousness. Alpha power was consolidated in posterior regions during baseline, 30- and 180-minutes post-ROR (Fig. 4D). An increase in frontal alpha power is visually apparent during the induction, unconscious, 30-, 10- and 5-minutes pre-ROR epochs. There was a small-to-medium, significant positive correlation between alpha power, and the frequency of motif 1 (R = 0.26, p < 0.001) and motif 5 (R = 0.22, p < 0.001).

### Network motifs reflect changes in long-range and short-range functional connections

The average total distances between nodes within a given motif was pooled across all participants and epochs and the distribution was plotted on a histogram (Figure 4, panel F). Nodes participating in motif 1 formed long-range connections (median total distance = 10.0 cm, range = 8.21 to 14.0 cm), while nodes participating in motif 5 formed short-range connections (median total distance = 4.21 cm, range = 2.91 to 8.55 cm). Nodes in motif 1 can either be connected to 1 other node (in the case of sources) or 2 other nodes (in the case of sinks) within the motif; individual connection distances therefore range between 4.11 and 14.0 cm in length. Nodes participating in motif 5 are always connected to 2 nodes; individual connection distances range between 1.45 and cm in length.

### Global network properties do not consistently distinguish between responsive and unresponsive states

We compared four global network properties (i.e. global efficiency, clustering coefficient, modularity and binary small-wordless) across the eight 5-minute epochs. We did not have sufficient evidence to conclude that global efficiency was significantly different from baseline at any time point according to Friedman’s test. A Bayesian repeated-measures ANOVA showed that the model including epoch as a factor was slightly more likely than the null model (*BF*_10_ = 14.985); however, there was anecdotal evidence for decreases in global efficiency from baseline during the unconscious epoch (*BF*_10,*U*_ = 1.349) and 5-minutes pre-ROR (*BF*_10,*U*_ = 1.112) (Fig. 5A). The clustering coefficient changed significantly across epochs (*χ*^2^(7) = 34.667, p < 0.001, W = 0.470). Specifically, this measure significantly increased from baseline at unconscious, 30-, 10-, and 5-minutes pre-ROR epochs (p < 0.01, Fig. 5B). Changes in binary small-worldness were driven by changes in the clustering coefficient; this metric was therefore also significant (*χ*^2^(7) = 34.481, p < 0.001, W = 0.448). Specifically, it was increased from baseline at the unconscious epochs, as well as 30-, 10-, and 5-minutes pre-ROR (p < 0.05, not plotted due to redundancy). Modularity changed significantly across epochs (*χ*^2^(7) = 32.481, p < 0.001, W = 0.277) and was significantly increased from baseline at 30-, and 5-minutes pre-ROR (p < 0.01, Fig. 5C). Only clustering coefficient and binary small-worldness statistically distinguished the unconscious epoch from baseline.

**Figure 5.**
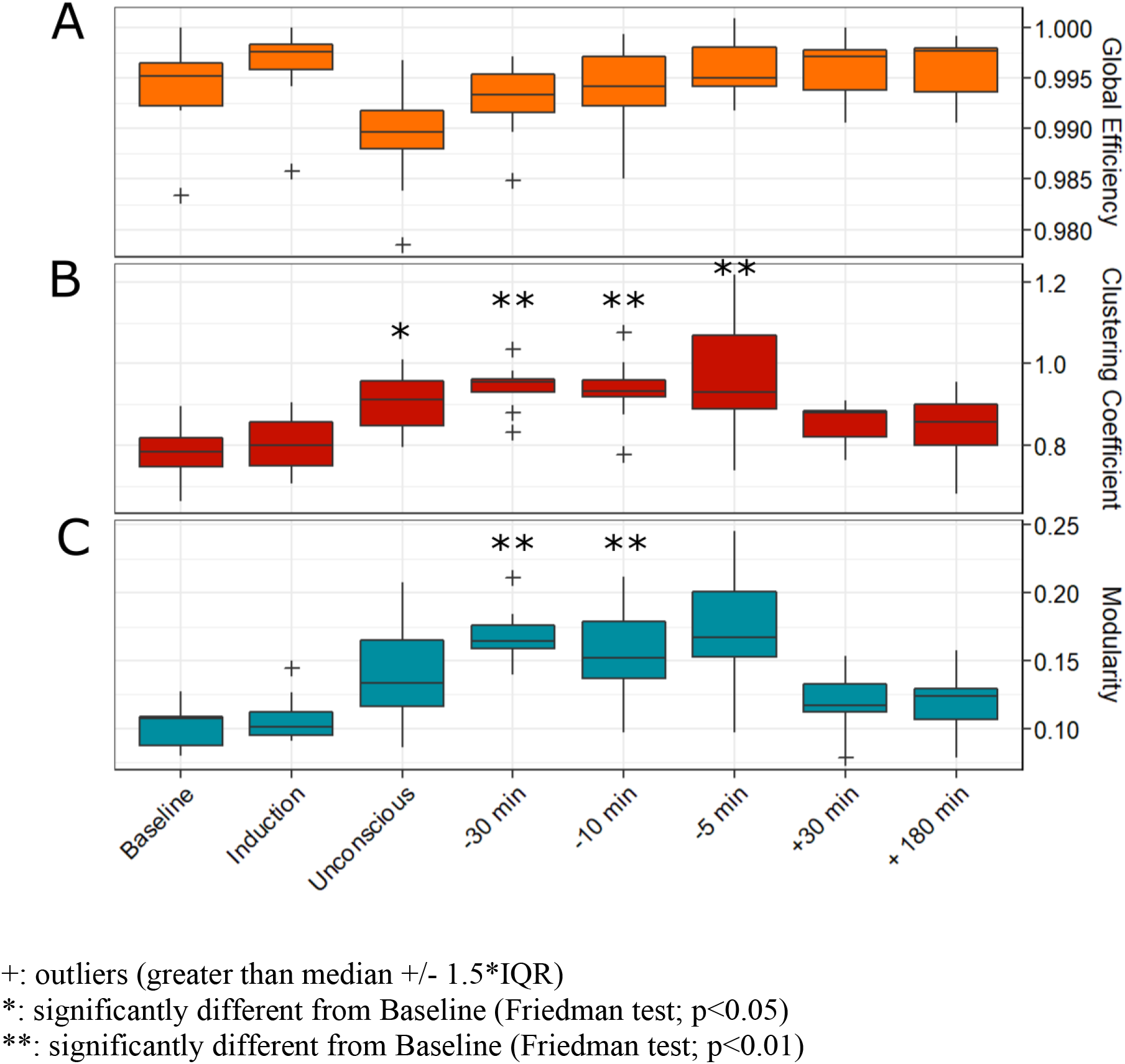
Brain network properties across the experimental period. The functional brain network of each participant was constructed using a binarized wPLI matrix. From this network, we calculated global graph theoretical network properties, including global efficiency (A), clustering coefficient (B), binary small-worldness and modularity (C), across the eight analysis epochs. Changes in binary small-worldness were driven by changes in the clustering coefficient; this metric was therefore also significantly increased from baseline at unsconsious, 30-, 10-, and 5-minutes pre-recovery of consciousness epochs (not plotted due to redundancy). Boxes represent the interquartile range, with the lower and upper limit of boxes indicating 1^st^ and 3^rd^ quartiles, respectively. The median is marked by the black horizontal line inside the boxes. The whiskers outside the boxes extend to the lowest and highest observations that are not outliers, while outliers are marked by a cross, and are defined as values greater than ±1.5 times the interquartile range.

### Network efficiency and clustering topology do not spatially reorganize across states of consciousness

To further explore our negative findings for global network properties, efficiency and clustering coefficient were plotted on a topographic head map at a nodal level (i.e. prior to averaging across all nodes to obtain global network measures). Unlike network motifs, we observed no consistent spatial organization of efficiency, or any reorganization of efficiency and clustering coefficient across states of consciousness, even at the individual subject level (see Supplemental Figure 5 for a sample individual participant).

## Discussion

We investigated motifs in human directed functional EEG networks to identify statistically significant motifs associated with states of consciousness, and to reveal the temporal changes in these basic network building blocks across loss and recovery of consciousness. The directed functional networks were constructed from high-density EEG data recorded from human participants before, during and after anesthetic-induced unconsciousness using a surgical levels of anesthesia, without the confounds of surgical stress, inflammatory burden or polypharmacy that often accompany anesthesia research. This protocol enabled us to assess consciousness-related transitions in brain networks. Networks were constructed from four epochs of behavioral responsiveness and four epochs of behavioral unresponsiveness. Three classes of 3-node motifs (i.e. ID = 1, 2 and 5) were significantly more likely to be embedded in these brain networks compared to null networks. These motifs were topologically distributed in distinct patterns that re-organized with changes in levels of consciousness. Motif 1 consisted of two independent nodes connected to a shared third node. This motif was constituted by long-range connections, with sources predominant in anterior and central regions during conscious wakefulness, and dominant in posterior regions during unresponsive states. The topological distribution of nodes participating in motif 1 significantly reorganized during all four unresponsive epochs. Motif 5 consisted of three interconnected nodes linked by short-range connections. Nodes participating in motif 5 were concentrated in posterior brain regions during baseline conscious wakefulness; anterior nodes increased their participation in this motif during the unconscious and 30-minutes pre-ROR epochs. The topologic distribution of motif 5 nodes returned to baseline patterns by 10-minutes pre-ROR, and maintained this pattern through all subsequent epochs.

It is instructive to examine the characteristics of the significant motifs in finer detail. These three motifs can be divided into two sub-types, according to their structure: chain-like motifs (ID = 1 and 2) and loop-like motifs (ID = 5). In the chain-like motifs, two nodes that are not directly linked are integrated through an intermediate third node. These chain-like motifs have been implicated in the communication between functional modules of the brain, both in humans and macaques [22,26,27]. Other studies have linked hub regions -- which play an important role in the information integration of the brain network -- to the intermediate apex of these chain-like motifs, and suggest that densely inter-connected hubs (the rich club) form a stable synchronization core through these chain-like motifs [28]. Our analysis of motif 1 is highly consistent with these previous observations. We demonstrated that motif 1 consists of long-range connections, with posterior sinks (i.e. the intermediate apex) during all responsive epochs and anterior sinks during all unresponsive epochs. This reconfiguration of modular architecture is consistent with, and provides a functional mechanism for, the anteriorization of network hubs observed during anesthetic-induced unconsciousness [29]. The significant topological disruption of this motif across all unresponsive epochs is also consistent with previous studies associating the disruption of long-range functional connectivity with loss of consciousness [16,30], lending credibility to our analysis. The loop-like motifs form a tight loop for information processing, enabling local integration to achieve functional specification. The loop-like motif in our analysis (ID = 5) consisted of short-range connections that were dominant in posterior regions in the baseline epoch, and shifted to an anterior-dominance during unconsciousness. The fact that this motif was no longer significantly distinct from its baseline pattern 10-minutes prior to the return of responsiveness suggests that the re-establishment of short-range connections may be necessary but not sufficient for behavioral responsiveness.

If consciousness is a pre-requisite for intentional behavior [31], then networks sustaining consciousness may return prior to the ability to understand and willingly respond to a command. The dissociation between consciousness and responsiveness has recently been established on the level of brain network dynamics, where network changes were linked to state of consciousness rather than behavioral responsiveness [32]. Our findings regarding the temporal course of motif topological reconfiguration may also suggest that distinct network processes underly consciousness and responsiveness. Although the topology of motifs 1 and 5 were spatially complementary and although they were both significantly disrupted during unconsciousness, motif 1 did not return to baseline patterns until ROR, while motif 5 returned at least 10 minutes before ROR. Motif 1 therefore appears to accompany behavioral responsiveness, and to be a marker of both the cognitive capacity to understand and respond to a command, and the motor ability to execute that command. In contrast, motif 5 returned prior to behavioral responsiveness, suggesting that it may constitute a prerequisite of consciousness. Moreover, Motif 5 may be a potential indicator of the imminent return to responsiveness, which would be of significant clinical value in monitoring for intraoperative awareness [33], and in assessing pathological unconsciousness such as unresponsive wakefulness syndrome and minimally conscious state [34].

We explored several other potential explanations regarding the mechanism driving the re-organization of network motifs across states of consciousness. First, we investigated the hypothesis that the observed motif re-organization was associated with anterior-posterior shifts in alpha power across states of consciousness. The anteriorization, or “frontal dominance” of alpha power has long been associated with anesthetic-induced unconsciousness [10,11,35,36], and the reorganization of motifs 1 and 5 exhibited a similar anterior-posterior shift in dominance across states of consciousness. There were small-to-medium correlations between alpha power and motif topology, indicating that the shift in alpha power may be a contributing factor to the observed shift in motif distribution. Alternatively, it is possible that both alpha power and motif topology are independent markers that reflect anesthetic-induced reconfiguration of network dynamics. Though the mechanisms of alpha-power anteriorization are still not well understood, they are posited to reflect alterations in cortico-thalamic interactions – essential for consciousness processing and cognitive function [37–40] – caused by the effect of anesthetic drugs on various thalamic nuclei [41,42]. Conversely, simulations with model complex networks have shown that nodes with larger degrees (e.g. network hubs) have larger amplitudes [15]. As the apex of motif 1 has also been co-located with network hubs [22], the correlation of alpha power and motif topology may simply reflect two epiphenomena markers of anesthetic-induced shifts in network hub location. Second, we investigated the hypothesis that changes in motif topology were driven by anesthetic-induced changes in motif frequency across the experiment. Anesthetic-induced unconsciousness has been associated with a decrease in the strength of directed functional connectivity across brain regions [15,43,44], which could potentially be accompanied by a decrease in motif frequency. However, the total frequency of each motif was not statistically different between any of the 8 epochs or across states of consciousness (conscious vs. unconscious), eliminating this as a plausible explanation for the observed changes in motif topologic distribution.

Our results highlight the value of network motifs as a complementary node-based measure to global network properties. Global network properties have proven useful in characterizing altered states of consciousness or changes in responsiveness [12–16,30,45–50]. Here, we showed that some global network properties were altered as of result of anesthesia, and recovered prior to or in parallel with the return of behavioral responsiveness. Similar to previous findings [12,51], we showed that unresponsive brain networks have increased clustering coefficients and modularity during some unresponsive epochs, though only clustering coefficients and binary small-worldness were statistically distinct from baseline during the unconscious epoch, and across all unresponsive epochs. Our Bayesian analysis suggests that the lack of statistically significant differences for global efficiency and modularity (during the unconscious epoch) is a result of our small sample size and the high variability between subjects. Unlike these global network properties, the topologies of motifs 1 and 5 were significantly different from baseline during the unconscious epoch (and other subsequent unresponsive epochs), in spite of the small sample size and high variability. These different results may suggest that macro-scale (i.e. global) vs. meso-scale (i.e. node-based) network properties reflect divergent information [52], and highlights the complementary information that motifs can provide in describing brain network correlates of consciousness.

The results of this study must be interpreted in light of several limitations. First, this was a small study of young healthy volunteers, and the patterns of motif distributions observed herein may not generalize to surgical patients of varying ages and comorbidities, nor to brain-injured patients. Secondly, loss and recovery of consciousness were indirectly assessed through behavioral responsiveness. Is it therefore impossible to confirm whether participants were truly unconscious during all epochs of unresponsiveness, or whether they retained awareness but had a loss of motor control or of the cognitive abilities necessary for understanding commands and initiating proper responses. Given the MAC of 1.3 during the 3-hour anesthetic period, we are confident that study participants were deeply anesthetized during the anesthetic protocol and during the “unconscious” epoch, which took place in the first five minutes following the end of isoflurane administration. However, unresponsiveness is not equivalent to unconsciousness [53], and awareness may still be present despite unresponsiveness. There have been accounts of intraoperative awareness and no method has yet proven to be completely effective in detecting consciousness (i.e. awareness) during an anesthetic state [54]. Thirdly, the “unconscious” epoch represents the first five minutes after isoflurane has been turned off, rather than during isoflurane administration. However, in these 5 minutes immediately after the 3-hour anesthetic period at surgical levels, participants were unresponsive, and remained completely unresponsive for over 30 minutes. This was a state of unresponsiveness where patients still had high levels of isoflurane concentration in their brain and blood, without the direct effect of isoflurane administration. Fourthly, as our directed functional network was constructed using directed phase lag index (dPLI), our analysis was limited to motifs with unidirectional connections between nodes. In the class of 3-node motifs, this restricted the scope of our analysis to 5 of the 13 potential 3-node motifs. While the patterns of motif distribution described in this paper remain valid, it is possible that the motifs highlighted herein are in fact a subgraph (i.e. a graph formed by a subset of the vertices of a larger graph) of bidirectional motifs. Constructing the directed functional network with a metric such as symbolic transfer entropy would enable non-complementary measures of bi-directional interactions between nodes, and merits future work [55]. Sixth, our interpretation of the topologic distribution of motif 1 as a marker of the recovery of responsiveness is limited by the study design, where the first available resting-state epoch post-recovery of responsiveness occurs 30 minutes after the return of responsiveness. Analysis of the networks constructed from epochs immediately following recovery of responsiveness are warranted to confirm the relationship of this motif distribution to an individual’s global state of consciousness [25]. Finally, our analysis was conducted on the level of the EEG sensors, which can record changes in brain activity that do not occur proximal to the electrodes under analysis; it is possible that the topographic changes in motifs reflect distant cortical interactions that are widely projected to many cortical sites. Future work should use source reconstruction or current source density analysis to model the data in source space to further illuminate the neurophysiological underpinnings of motif configurations and their causal relationship to states of consciousness.

This study provides preliminary evidence that changes in states of consciousness induce a structured re-organization in the topology of functional network motifs. Though motif frequency remains constant under anesthetic-induced unconsciousness, the motif topology associated with conscious wakefulness shifts during unconsciousness and returns either prior to, or in parallel with the recovery of responsiveness, in a motif-specific manner. As such, motifs could improve the monitoring, identification and prognostication of consciousness. As nodal properties of functional networks, motifs may complement global network properties in the study of the neural correlates of consciousness.

## Methods

### Participants

Nine healthy volunteers (5 men; 24.4 ± 1.0 years old) were recruited at the University of Michigan, as part of the Reconstructing Consciousness and Cognition study (NCT01911195) [56]. A tenth participant was initially recruited but was subsequently excluded from all analyses due to excessive noise artifacts in the EEG. The study was approved by the Institutional Review Board of the University of Michigan (HUM0071578). All methods were performed in accordance with relevant guidelines and regulations, and written informed consent was obtained from all participants. As this was an observational study, it was not registered in a clinical trial registry. Participants were included if they were between 20 and 40 years old, had a BMI < 30 kg/m^2^, satisfied the criteria for the American Society of Anesthesiologists Physical status I or II [57], had an easily visualized uvula and were able to provide signed informed consent. Participants were excluded if they had physical indication of a difficult airway, family history of problems with anesthesia, obstructive sleep apnea, neuropsychiatric disorders, hypertension, cardiovascular disease, reflux, sleep disorders, postoperative nausea or vomiting, motion sickness, or reactive airway disease. Participants were also excluded for pregnancy, past or current use of psychotropic medications, current tobacco or alcohol use exceeding 2 drinks/day, positive urine toxicology test, allergy to eggs, egg products, or soy.

### Anesthetic protocol

Participants underwent a 3-hour anesthesia protocol at surgical levels. As previously described [56], participants underwent a standard clinical preoperative history and physical examination on the day of the study. The anesthetic protocol took place in an operating room. Standard electrocardiogram, non-invasive blood pressure cuff, pulse oximeter, and capnography were used for constant monitoring during the protocol. Patients were pre-oxygenated by face mask prior to induction of general anesthesia with a stepwise increasing infusion rate of propofol: 100 mcg/kg/min × 5 minutes increasing to 200 mcg/kg/min × 5 minutes, and then to 300 mcg/kg/min × 5 minutes. After 15 minutes of propofol administration, inhalation of 1.3 age-adjusted minimum alveolar concentration (MAC) of isoflurane was started [58]. Loss of consciousness occurs generally around 0.3 MAC [59], 1.0 MAC can abolish evoked related potentials [60], and 1.3 MAC produces suppression of the sympathetic nervous system [61,62]. Consequently, 1.3 MAC reflects surgical anesthesia and is deemed a deep anesthetic state [62]. A laryngeal mask was inserted orally, a nasopharyngeal temperature probe was placed, and the propofol infusion was discontinued. Anesthetized subjects received 1.3 age-adjusted MAC inhaled isoflurane anesthesia for 3 hours. Blood pressure was maintained within 20% of baseline pre-induction values using a phenylephrine infusion or intermittent boluses of ephedrine, as necessary. To prevent post-anesthetic nausea and vomiting, participants received 4 mg of intravenous ondansetron 30 minutes prior to discontinuation of isoflurane.

At the end of the 3-hour anesthetic period, isoflurane was discontinued and we started an audio loop command that was played every 30 seconds, asking the participant to squeeze their left or right hand twice (randomized order). Recovery of consciousness was estimated through return of responsiveness, which was defined as the earliest instance in which participants correctly responded to two consecutive audio loop commands. The laryngeal mask was removed when deemed medically safe by the attending anesthesiologists.

### Electroencephalographic acquisition and preprocessing

EEG was acquired using a 128-channel system from Electrical Geodesics, Inc. (Eugene, OR) with all channels referenced to the vertex (Cz). Electrode impedance was maintained below 50 kΩ prior to data collection and data were sampled at 500 Hz. Throughout the experiment, data were visually monitored by a trained investigator to ensure continued signal integrity. All data preprocessing was performed in EEGLAB [63]. Five-minute epochs of EEG were extracted for the following time points: 1) baseline; 2) induction; 3) unconscious; 4) 30 minutes prior to recovery of responsiveness (ROR); 5) 10 minutes prior to ROR; 6) 5 minutes prior to ROR; 7) 30 minutes post-ROR; and 8) 180 minutes post-ROR (Fig. 1). Four of these epochs were associated with a state of responsiveness (epochs 1, 2, 7, 8) and four were associated with a state of unresponsiveness (epochs 3, 4, 5, 6). EEG data were bandpass filtered between 0.1 and 50 Hz and re-referenced to an average reference. Non-scalp channels were discarded, leaving 99 channels for the subsequent analyses. All epochs and channels with noise or non-physiological artifacts were visually identified and removed. We then band-pass filtered the cleaned EEG data into delta (1–4 Hz), theta (4–8Hz), alpha (8–13Hz), and beta (13–30 Hz) frequency bands. Analysis of the alpha band were presented in the main body of the paper, while analyses of other frequency bands are included in the supplementary material.

### Functional Connectivity

To construct a functional brain network from EEG data, we used the weighted phase lag index (wPLI) [64], and the directed phase lag index (dPLI) [65] (Fig. 6), which are robust methods to avoid volume conduction confounds in the data [66]. This was done using custom MATLAB scripts (version R2018b).

**Figure 6.**
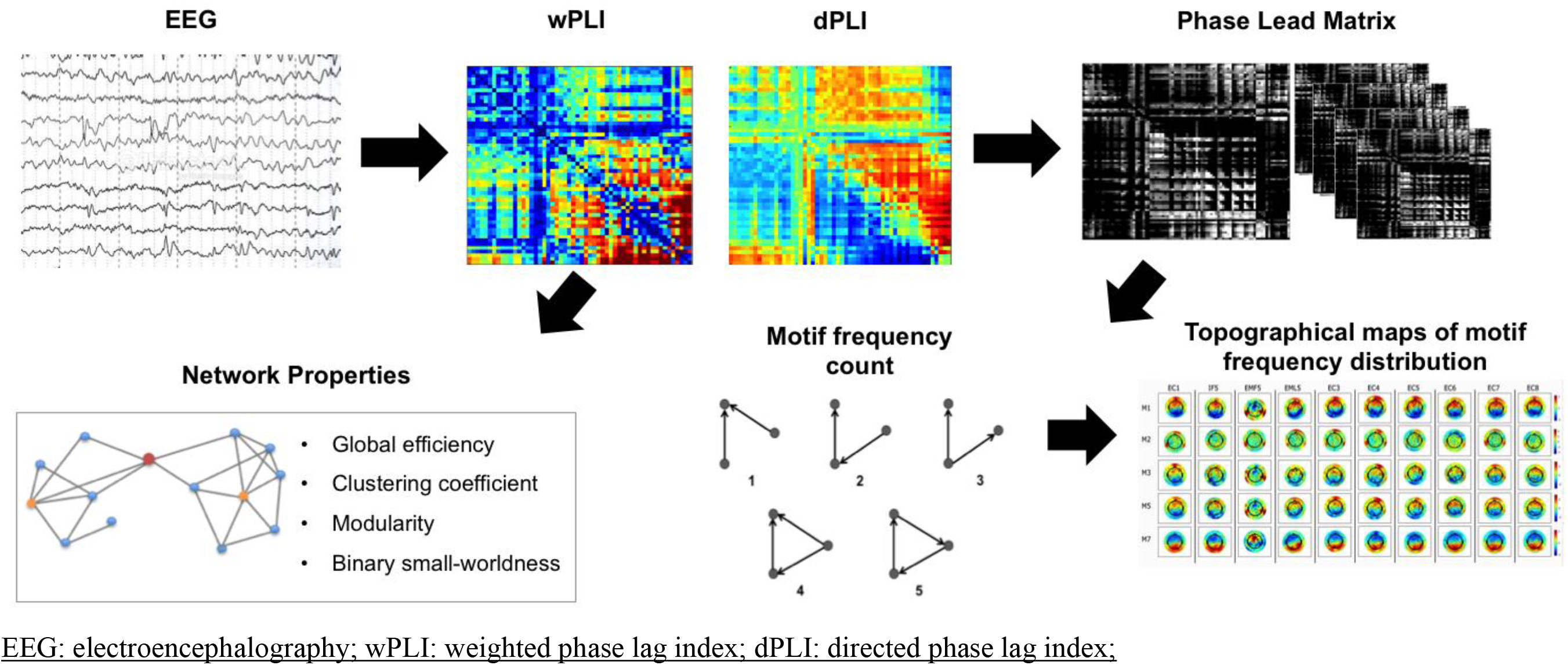
Pipeline for extracting network properties and motifs EEG time series. Each 5-min EEG epoch was used to construct a wPLI and dPLI matrix. Network properties were then calculated based on the wPLI matrix, while the dPLI matrix was used to create a phase-lead matrix, which set all non-phase-leading values to 0 (black), and normalized the remaining non-zero values between 0 and 1. Motifs were then extracted from the phase lead matrix, and their frequency (i.e frequency of participation of a node in each of the possible types of motifs) and topology (i.e. topological distribution of participating nodes in each motifs) were calculated.

First, we segmented the cleaned EEG data into delta (1-4 Hz), theta (4–8Hz), alpha (8– 13Hz), and beta (13-30 Hz) frequency bands using band-pass filtering methods (Butterworth filter). wPLI was calculated using the following formula:

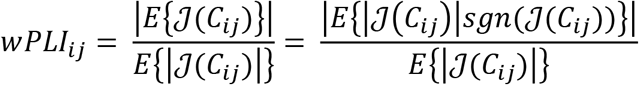

where ℐ(*C_ij_*) is the imaginary part of cross-spectrum *C_ij_* between signals *i* and *j* [64]. The cross-spectrum *C_ij_* is defined as Z_*i*_Z_*j*\*_, where Z_*i*_ is the complex value Fourier spectra of the signal i for each frequency, and Z_*j*_* is the complex conjugate of Z_*j*_. *C_ij_* can be written as *Re^iθ^*, where R is magnitude and *θ* is the relative phase between signal *i* and *j* [64]. A wPLI value of 1 indicates complete phase locking between the two signals (i.e. that the instantaneous phase of one signal is leading the other). Conversely, a wPLI value of 0 indicates no consistent phase-lead or -lag relationship. To know the direction of the phase-lead/phase-lag relationship between channels *i* and *j* in the wPLI matrix, we calculated the dPLI [65]. First, the instantaneous phase of each EEG channel was extracted using a Hilbert transform. The phase difference *Δφ_t_* between all the channels was then calculated where *Δφ_t_* = *φ_i,t_* − *φ_j,t_*, t = 1,2,…,N, where N is the number of samples in one epoch, and *i* and *j* include all channels. dPLI was then calculated using the following formula:

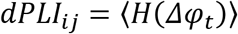

where H(x) represents the Heaviside step function, where H(x) = 1 if x > 0, H(x) = 0.5 if x = 0 and H(x) = 0 otherwise. Thus, if on average signal *i* leads signals *j*, dPLI will be between 0.5 and 1, and if signal *j* leads signal *i*, dPLI will be between 0 and 0.5. If there is no phase-lead/phase-lag relationship between signals, dPLI = 0.5.

For both the wPLI and dPLI matrices, we controlled for noise-induced phase relationships using surrogate datasets in which we randomized the phase relationship between two channels, while maintaining their spectral properties. More specifically, using the instantaneous phase for each channel pair *i* and *j*, we maintained the phase time series for *i* and scrambled the time series for *j* from 0 to x, by swapping it for the time series from x to n, where n is the number of samples in one epoch, and 0 < x < n. Data segments used to generate surrogate wPLI/dPLI matrices were 10 seconds in length, and each was permuted 20 times to generate a distribution of values representing the spurious connectivity. The wPLI and dPLI values of the original, non-shuffled EEG data were compared to this distribution of surrogate data using a Wilcoxon signed rank test and were set to 0 (wPLI) or 0.5 (dPLI) if they did not achieve statistical significance. Statistical significance was set to *p* < 0.05.

### Network motifs

In a directed network, a motif is a subnetwork consisting of N nodes and at least (N-1) edges linking the nodes in a path [18,19]. In this study, we investigated network motifs of N = 3 using a unidirectional network, for which 5 motifs are possible (Fig. 2). Each EEG electrode represented a single node; a 3-node motif therefore represents a specific pattern of interconnection between any three electrodes on the scalp (i.e. motifs are not constrained to connections between neighboring electrodes). While motifs comprised of other numbers of nodes are possible, we restrict the analysis in this paper to N = 3 due to the exponential increase in computational complexity that co-occurs with increasingly higher numbers of nodes.

Network motifs were computed using custom MATLAB scripts and the BCT. Prior to computing the network motifs, we normalized the dPLI matrix into a non-symmetrical phase-lead matrix, in order to remove redundant information. Given the nature of the dPLI matrix, a value at position (i,j) is necessarily the opposite of a value at position (j,i). We therefore set any value that was below 0.5 in the dPLI matrix, corresponding to a non-phase-leading value, to 0. The remaining non-zero values were normalized between 0 and 1. This normalized phase-lead matrix was then used to calculate the frequency of participation of each node in each motif [67].

Two levels of surrogate analysis were used to calculate whether the probability of occurrence of motifs is beyond chance (i.e. whether a motif is significantly present). First, we corrected for the surrogate number of motifs calculated at each node as follows: using BCT [67], we generated 100 null networks that preserved the degree and strength distribution from the normalized phase lead matrix. For each of the 100 null networks, we calculated the mean frequency of each motif, for each node. Second, we corrected for the surrogate number of motifs across each network: for each motif, we calculated the total aggregate frequency of motifs in the network by summing across all nodes. We then calculated a network Z-score per motif against the distribution of the total motifs in the 100 null networks as follows:

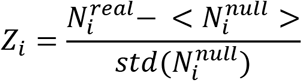

where 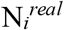 is the aggregate frequency of motif *i* across the network and 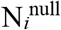 is the aggregate motif frequency of the 100 null networks [22]. If *Z_i_* was greater than 1.96, the motif was considered to be significant in the network; if not, it was considered non-significant and set to a frequency of 0.

### Analysis of global network properties

The functional brain network was constructed using the wPLI of all pairwise combinations of electrode channels. We constructed a binary adjacency matrix A_*ij*_ using a threshold of 35%: if the wPLI_*ij*_ value of nodes *i* and *j* was within the top 35% of all wPLI values, A_*ij*_ = 1; otherwise, A_*ij*_ = 0. We chose a binarized approach to be coherent with the majority of studies assessing global network properties, which have also used a binarized approach to construct their graph [12,15,23,29,68–70]. The 35% threshold was selected because it was previously shown to be the optimal threshold to avoid an isolated node in the EEG network during baseline [70]. From the binary adjacency matrix, we calculated global network properties using the Brain Connectivity Toolbox (BCT) [67], including global efficiency, clustering coefficient, and modularity (Fig. 6). Global efficiency is the inverse of the average shortest path length 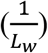, where Lw is the average of the shortest path lengths (L_*ij*_) between all pairs of nodes in the network [48]. The clustering coefficient, calculated by averaging the clustering coefficients of all individual nodes (C_*i*_), represents the degree to which nodes of a graph tend to cluster together, such that higher values imply networks with highly clustered or regular structures [71]. The modularity of the network represents the strength of division of a network into modules, such that high modularity implies a network with strong within-module connections and weak between-module connections [72]. Modularity was calculated using the Louvain algorithm. Finally, small-world organization is characterized by high levels of clustering and short path lengths, meaning that all nodes of a network are linked through relatively few intermediate steps, though they are only directly connected to a few, mainly neighboring, nodes [1]. Binary small-worldness is therefore the ratio of the clustering coefficient to the average path length, after both metrics are normalized against random networks [73]. Global efficiency and clustering coefficient were normalized by taking the ratio between the true network metric and the average metrics computed for 10 random binary networks, which were generated by shuffling the empirically-generated network’s edges 10 times while preserving the degree distributions, as previously done [18,74].

### Topological configuration of network motifs across states of anesthetic-induced unconsciousness

The distribution of nodes participating in each motif was visualized on a topographic head map, using the *topoplot* function of EEGLAB [63]. Changes in the topologic distribution of participating nodes were assessed using cosine similarity on the topographic head maps, defined as:

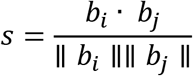

where b_*i*_ and b_*j*_ are vectors containing motif frequencies for states *i* and *j* [23]. Cosine similarity ranges from −1 to 1, where 1 indicates identical topological distribution, −1 indicates completely opposite topological distribution, and 0 indicates orthogonality or decorrelation. This measure was computing using custom MATLAB scripts.

### *Post hoc* analyses

Once significant motifs were identified, we conducted two additional analyses to further interpret our results.

#### Topological distribution of source nodes within a motif

Using the dPLI, we conducted a source node analysis, in which we assessed the lead-lag relationships in every motif to identify which nodes were origins of information flow (i.e. “sources”), and which were destinations of information (i.e. “sinks”). Nodes that were phase-leading other nodes within a motif on their phase relationship were considered sources. Across the total frequency of participation of each node in a given motif, the total number of times that the node behaved as a source was calculated. To assess the topological distribution of sources within the network, the source total for each node was normalized by computing its Z-score relative to all other node source totals in a given network. These Z-scores were then plotted on a topographic map to visualize the relative distribution of sources within the network. This analysis was performed using the BCT and custom MATLAB scripts.

#### Topological distribution of alpha power

We investigated the topological distribution of alpha power as a possible explanation for the changes in anterior-posterior dominance observed in motifs across states of consciousness. The Pearson correlation between the motif frequency and the raw alpha power (*μ*V/Hz^2^) for each node in the network was calculated. Alpha power was averaged across 10-second windows and computed using a multi-taper power spectral density estimate (number of tapers = 3, time-bandwidth product = 2, spectrum window size = 3 seconds) from the Chronux package [75,76]. Correlation between alpha power and the frequency of participation of each node within a was visualized using a scatterplot, and quantified by computing the Pearson’s correlation (R) between x and y, where x is the motif frequency and y is the power for a given node, across all participants, time points and nodes, constructed using custom MATLAB scripts.

#### Distance distribution of motif connections

For each node *u*, we summed the Euclidean distance (*d*) between *u* and all of its connected nodes *v_i_* within a given motif, where *i* = {1,2} (i.e. up to two connected nodes). We repeated this for every motif *j* in the network, and then normalized by the motif frequency for that node (*f_u_*). Thus, for each node, we obtain the total distance between a node and all its connections within a motif, averaged across all the motifs that node participates in, according to the following formula:

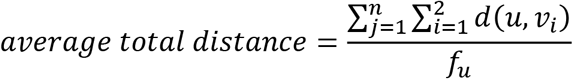

where *d*(*u*, *v_i_*) is the Euclidian distance between points *u* and *v_i_*.

We pooled the average total distances for motifs 1 and 5 across all participants and epochs and plotted their distribution using a histogram. To obtain individual connection distances, we divide the average total distance by the number of connections a node has.

### Statistical analyses

Due to our small sample size and high variability between participants, total motif frequency (i.e. the sum of motif frequency across all nodes), cosine similarity of motif topologies relative to baseline, and global network properties (i.e. global efficiency, clustering coefficient, binary-small worldness and modularity) were compared across epochs using a non-parametric Friedman’s test. Effect size was measured using Kendall’s coefficient of concordance (W). When Friedman’s test was significant, *post hoc* Bonferroni-corrected Conover’s tests were performed, comparing all variables between baseline and all other epochs (i.e. 7 comparisons). All tests were two-tailed with statistical significance set to *p* < 0.05.

When negative findings were observed for Friedman’s test, a Bayesian repeated-measures ANOVA was performed to investigate whether these negative findings were meaningful. Negative findings were considered to not be meaningful when the Bayes factor was greater than 1 (*BF*_10_, where 0 is the null model and 1 is the model including epoch as a factor).

Statistical analyses were conducted using JASP (version 0.12.2.0)

## Acknowledgements

This work was funded by the James S. McDonnell Foundation, St. Louis, MO (GM, MK and MA); the Canadian Institute for Health Research (FRN 152562, CD; Fredrick Banting and Charles Best Canada Graduate Scholarship – Masters, DN); the Fonds de Recherche du Québec – Nature et technologies (YM); the Fonds de Recherche du Québec – Santé (DN); and the Natural Science and Engineering Research Council of Canada (Discovery Grant RGPIN-2016-03817; SBM).

## Author Contributions

MA, MK, and GM conceived the study; VT, PP, GV, GG, EJ and SBM collected the experimental data; YM, DN, CD, and SBM analyzed the experimental data; CD, YM, DN, and SBM interpreted the findings; CD and SBM wrote the manuscript; DN and GM constructively reviewed the manuscript; all authors reviewed and approved the final version of the manuscript.

## Additional Information

Supplementary information accompanies this paper at https://doi.org/10.1038/s41598-019-41274-. All scripts used for network construction and analysis can be found at https://github.com/BIAPT/Scripts/tree/master/Motif%20Analysis%20Augmented%20(Source%2CTarget%20and%20Distance).

