## Supplementary Material for "Brain network motifs are markers of loss and recovery of consciousness"

### Appendix

### Brain network motifs are markers of loss and recovery of consciousness

#### Supplemental Figure 1

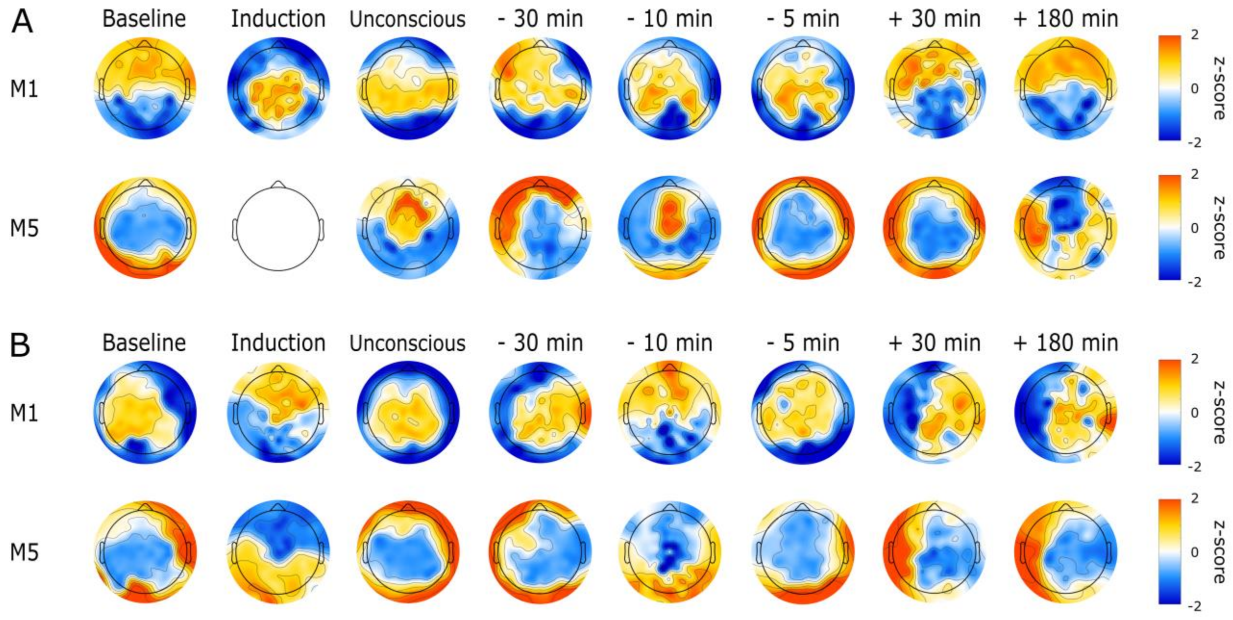


Topography of motif frequency in the alpha band (8–13 Hz) across states of consciousness. Visually compelling examples of motifs 1 and 5 (evident in 7/9 participants) are shown in panel A, while examples of non-compelling examples (evident in 2/9 participants) are in panel B. While the specific motif frequency distributions are different between individuals, motif 1 is central- or anterior-dominant during consciousness and shifts posteriorly during unconsciousness, while motif 5 is posterior-dominant during responsive states and involves anterior electrodes during unresponsive states.

**Supplemental Figure 2**

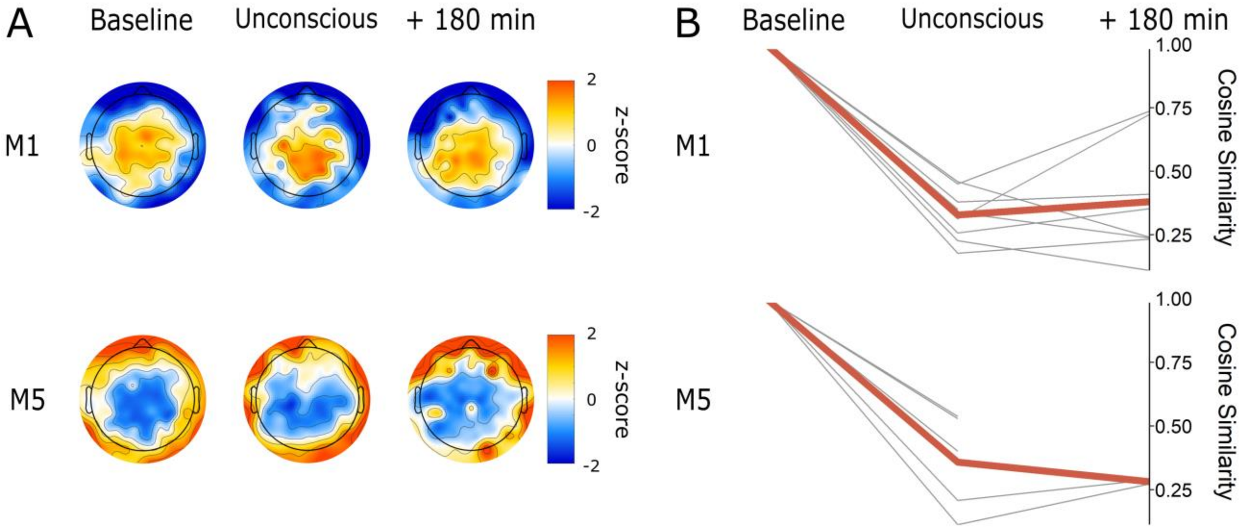


Motif frequency averaged across participants in the delta band (1–4 Hz). (A) Topographic maps indicate z-normalized motif frequency. (B) Cosine similarity between motif topographies at baseline, unconsciousness and 180-minutes post-return of responsiveness. Thin lines represent individual participants, while the thicker line represents cosine similarity averaged across participants. A reduced number of thin lines (< 9) indicates that motifs were non-significant for some participants. Here, cosine similarity shows that while motif topography shifts during unconsciousness, it does not return to its baseline state upon recovery of behavioral responsiveness. *M1 = motif 1, M5 = motif 5.*

**Supplemental Figure 3**

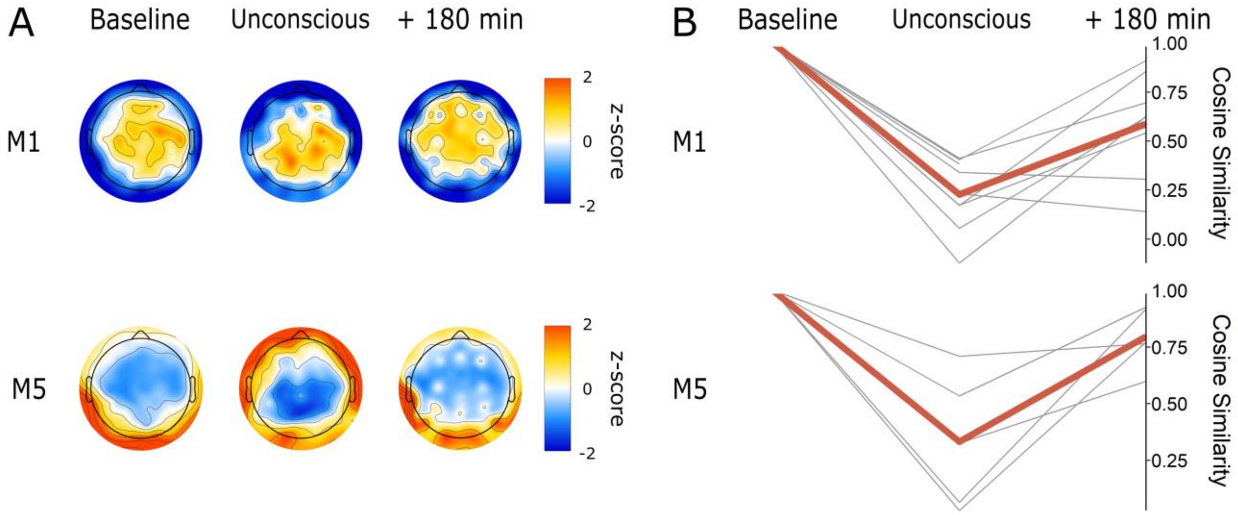


Motif frequency averaged across participants in the theta band (4–8 Hz). (A) Topographic maps indicate z-normalized motif frequency. (B) Cosine similarity between motif topographies at baseline, unconsciousness and 180-minutes post-return of responsiveness. Thin lines represent individual participants, while the thicker line represents cosine similarity averaged across participants. Cosine similarity supports what can be observed visually: motif 1 shifts slightly posteriorly while motif 5 shifts anteriorly during unconsciousness. Upon recovery of behavioral responsiveness, both motifs return to their baseline topography. *M1 = motif 1, M5 = motif 5.*

**Supplemental Figure 4**

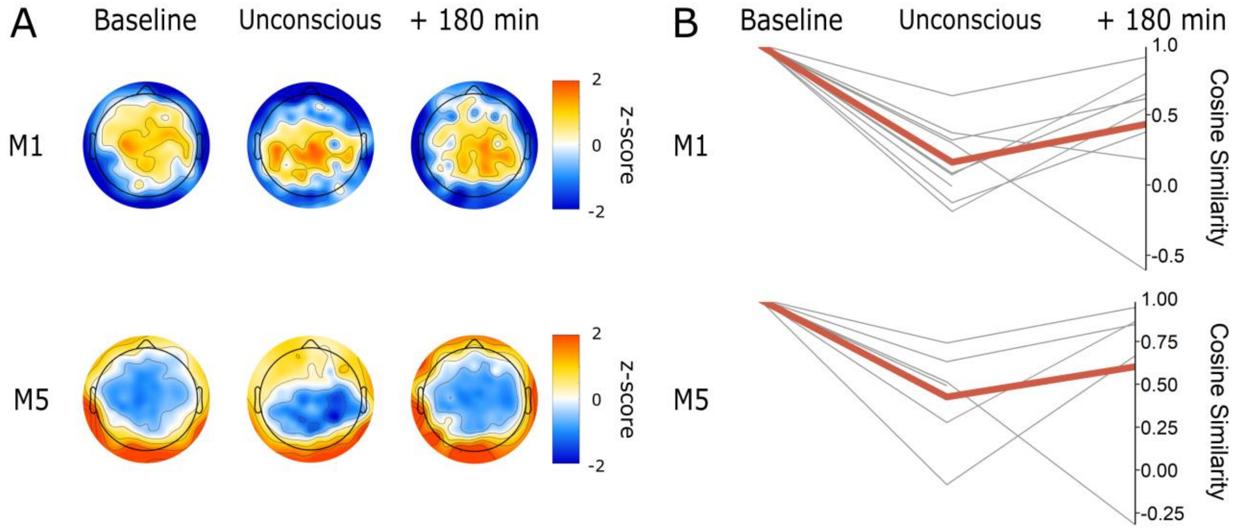


Motif frequency averaged across participants in the beta band (13–30 Hz). (A) Topographic maps indicate z-normalized motif frequency. (B) Cosine similarity between motif topographies at baseline, unconsciousness and 180-minutes post-return of responsiveness. Thin lines represent individual participants, while the thicker line represents cosine similarity averaged across participants. A reduced number of thin lines (< 9) indicates that motifs were non-significant for some participants. Here, cosine similarity shows that while motif topography shifts during unconsciousness, it does not return to its baseline state upon recovery of behavioral responsiveness. *M1 = motif 1, M5 = motif 5.*

**Supplemental Figure 5**

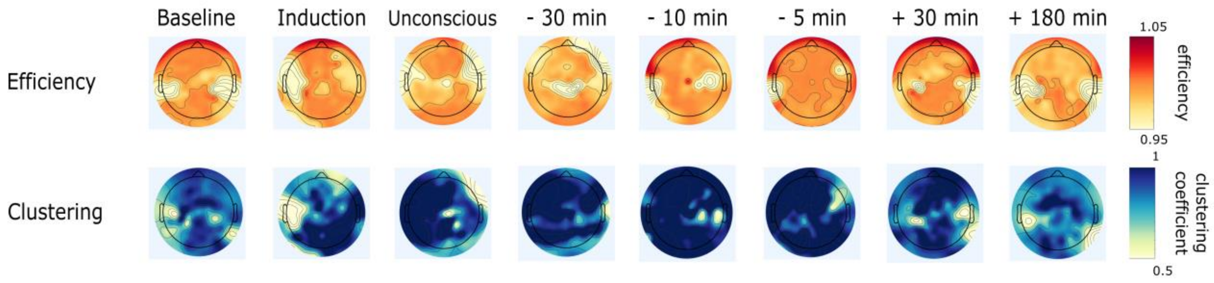


Topography of efficiency and clustering coefficient in the alpha band (8–13 Hz) across states of consciousness for two individual participants. The same participant as in Supplemental Figure 1 Panel A Motif 1 was used, to allow for comparison to their individual Motif 1 topography. Efficiency represents the inverse of the average path length between each node and all other nodes in the network. Clustering represents the clustering coefficient at each node. Both metrics were corrected against 10 random networks. There is no clear topographic organization of these metrics at baseline, or across epochs. We can observe a slight decrease in the magnitude of efficiency and an increase in the magnitude of clustering, which is in line with what we observe when analyzing the global efficiency and clustering coefficient (averaged across all nodes).
